# Disrupted cholinergic modulation can underlie abnormal gamma rhythms in schizophrenia and auditory hallucination

**DOI:** 10.1101/072504

**Authors:** Jung Hoon Lee

## Abstract

The pathophysiology of auditory hallucination, a common symptom of schizophrenia, has yet been understood, but during auditory hallucination, primary auditory cortex (A1) shows paradoxical responses. When auditory stimuli are absent, A1 becomes hyperactive, while A1 responses to auditory stimuli are reduced. Such activation pattern of A1 responses during auditory hallucination is consistent with aberrant gamma rhythms in schizophrenia observed during auditory tasks, raising the possibility that the pathology underlying abnormal gamma rhythms can account for auditory hallucination. Moreover, A1 receives top-down signals in the gamma frequency band from an adjacent association area (Par2), and cholinergic modulation regulates interactions between A1 and Par2. In this study, we utilized a computational model of A1 to ask if disrupted cholinergic modulation could underlie abnormal gamma rhythms in schizophrenia. Furthermore, based on our simulation results, we propose potential pathology by which A1 can directly contribute to auditory hallucination.

## 1. Introduction

Auditory hallucination is one of the most common symptoms of schizophrenia. The underlying pathophysiology of auditory hallucination remains elusive, but a line of studies proposed that primary auditory cortex (A1) may be involved generating auditory hallucination (K. Kompus, Westerhausen, & Hugdah, 2011; Kristiina Kompus, Falkenberg, Bless, Johnsen, & Kroken, 2013; F. Waters et al., 2012). They suggest that A1 is not a single source but a part of aberrant network responsible for auditory hallucination (Curcic-Blake et al., 2017; Powers III, Kelley, & Corlett, 2016). In addition, aberrant functional and anatomical connections between the left and right A1 (Christoph Mulert et al., 2012) and between A1 and other brain regions such as frontal lobes (Ford & Mathalon, 2005) were observed during auditory hallucination.

The studies mentioned above raise the possibility that A1’s neural circuits regulating endogenous inputs from other brain regions can underlie auditory hallucination. During auditory hallucination, A1 activation appears abnormal: its responses to auditory stimuli are reduced, but the spontaneous activity is enhanced (K. Kompus et al., 2011; Kristiina Kompus et al., 2013). This paradoxical activation pattern during auditory hallucination is consistent with abnormal gamma frequency band power in electroencephalography (EEG) observed in schizophrenia patients during auditory tasks (Hirano et al., 2015; Kwon et al., 1999; Spencer, 2009, 2011; Vierling-Claassen, Siekmeier, Stufflebeam, & Kopell, 2008), which suggests that we can infer the pathophysiology of auditory hallucination from abnormal gamma rhythms in schizophrenia. To address this possibility, we used a computational model of A1. Inspired by recent physiological findings (Roopun et al., 2010) that A1 receives top-down signals in the gamma frequency band (top-down gamma rhythms) from an adjacent association area (Par2) and that cholinergic modulation regulates A1 responses to top-down signals, we hypothesized that disrupted cholinergic modulation may induce abnormal gamma rhythms in schizophrenia. Thus, we first asked if disrupted cholinergic modulation could account for abnormal gamma rhythms during auditory tasks. We then studied the potential mechanisms by which A1 generates outputs, when high order auditory/cognitive areas perceive real auditory stimuli despite the absence of real sound (Kristiina Kompus et al., 2013; F. A. Waters, Badcock, Michie, & Maybery, 2006).

Our model A1 consists of pyramidal (Pyr), fast-spiking (FS) and non-FS interneurons; non-FS cells model low-threshold spiking interneurons which are known to express somatostatin; in fact, we focus on superficial layers of A1 projecting auditory signals to high order auditory areas (Douglas & Martin, 2004; Felleman & Van Essen, 1991; Markov & Kennedy, 2013). In the model, the pathological condition is simulated by lowering the excitability of non-FS cells since non-FS cells are known to have cholinergic receptors (Couey et al., 2007; Roopun et al., 2010; Xiang, Huguenard, & Prince, 1998). Our simulation results support our hypothesis that disrupted cholinergic modulation enhances baseline gamma rhythms and reduces stimulus-evoked gamma rhythms. More importantly, we note the two possible mechanisms by which A1 generates bottom-up signals even without stimulus inputs.

First, top-down gamma rhythms can make A1 Pyr cells fire in the pathological condition. As A1 outputs are independent from stimulus inputs, they make false auditory signals propagate to high order cognitive/auditory areas. Second, asynchronous top-down signals targeting FS cells can suppress A1 responses to the stimulus inputs. That is, FS cells can mediate corollary discharge, preventing A1 from responding to self-generated sound. In the model, when the strength of connections from a high order area to A1 is weakened, A1 responses to (simulated) self-generated sound grow stronger. This is consistent with the hypothesis that the failure of corollary discharge causes auditory hallucination, which is supported by the reduced functional connectivity from frontal to temporal lobes (Ford & Mathalon, 2005) in people with schizophrenia.

## 2. Methods

We used the peer reviewed simulator named NEST (Gewaltig & Diesmann, 2007) to build a network model. All neuron models and synapse models are natively supported by NEST. As shown in Figure 1A, we implemented superficial layers of A1 and three external populations. Specifically, A1 consisting of 400 Pyr, 70 FS and 30 non-FS cells interacts with three external populations of 100 Pyr cells (ovals in Figure 1A). The ratio of cell types are based on 1) the ratio of excitatory and inhibitory cells, which is 4:1, and 2) the fraction of somatostatin-positive cells, which is 30% (Rudy, Fishell, Lee, & Hjerling-Leffler, 2011); our non-FS cells model low-threshold spiking interneurons that express somatostatin.

**Figure 1:**
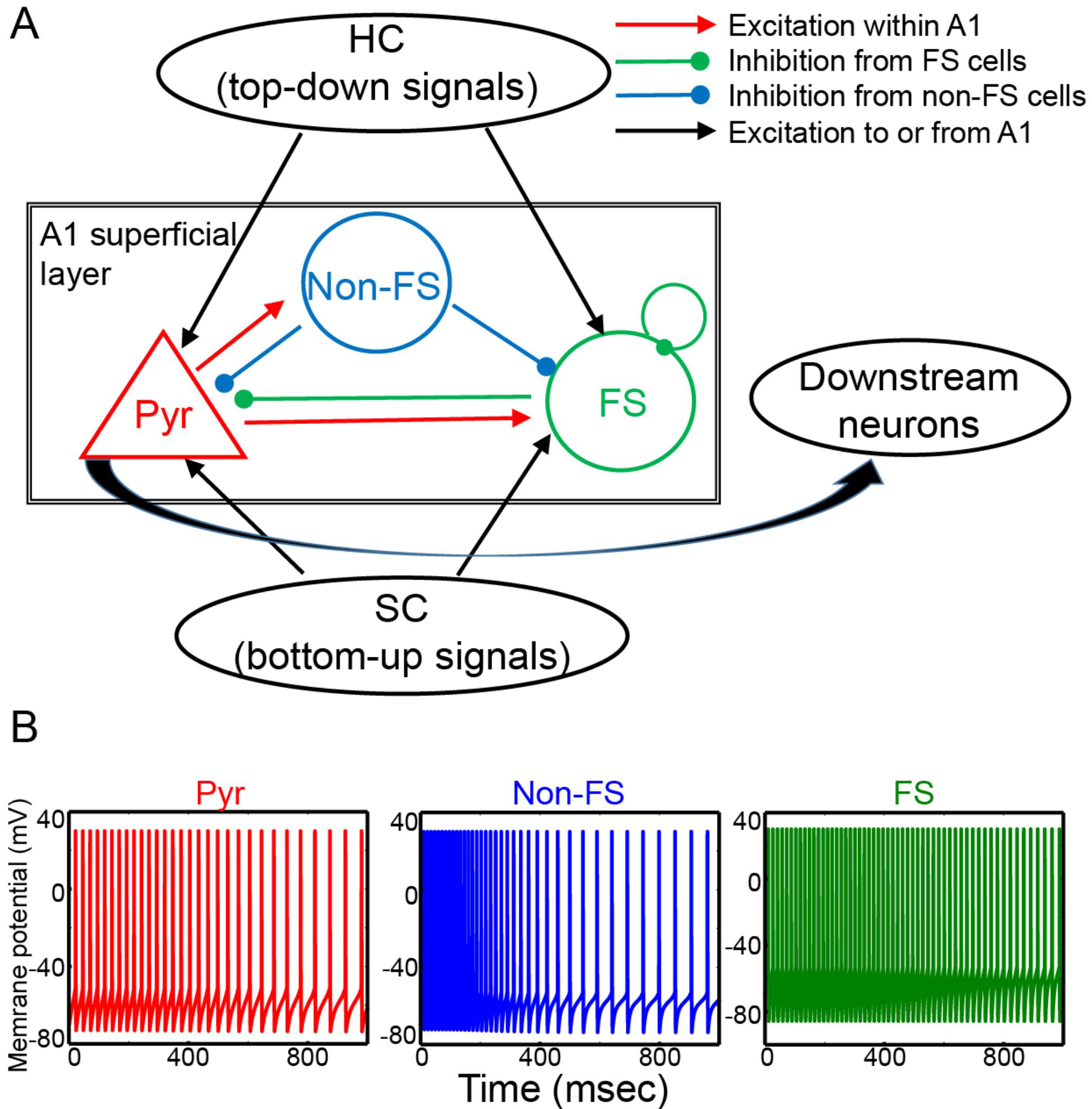
The structure of the model. **(A)**, The superficial layer of A1 is explicitly modeled using three cell types (Pyr, FS and non-FS cells in red, blue and green, respectively). In the model, there are three external populations (shown as ovals) of Pyr cells interacting with A1. The first two populations (HC and SC) project top-down and bottom-up gamma rhythms into A1. The last population (downstream neurons) receives synaptic inputs from A1 Pyr cells. **(B)**, The firing patterns of the three cell types in response to 20 pA tonic current injection.

The three external populations, each of which includes 100 cells, represent cortical areas interacting with A1. HC, SC and downstream neurons model high order cognitive areas such as an association cortex, subcortical areas and high order auditory cortices to which A1 projects its outputs, respectively. Downstream neurons passively receive inputs from A1, while HC and SC provide synaptic inputs to Pyr and FS cells in A1. They generate either synchronous and asynchronous inputs. The synchronous outputs are generated via inhomogeneous Poisson process, which is realized by the NEST’s native device named ‘sinusoidal Poisson generator’. Specifically, the firing rate (*R*) of the Poisson spike trains evolves over time in accordance with Equation 1:

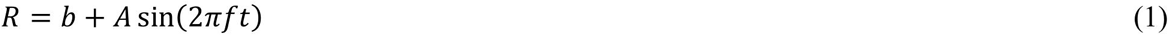

where *f* (frequency) represents the frequency of *R*; where *A* and *b* determine the peak firing rate and baseline, respectively. The synchronous Poisson spike trains are generated by HC and SC neurons with parameters listed in Table 1.

**Table 1:**
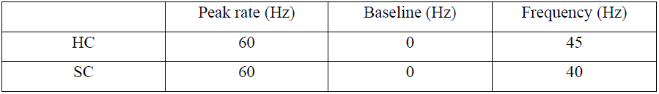
Parameters for sinusoidal Poisson generator. This NEST-native device (Gewaltig & Diesmann, 2007) generates inhomogeneous oscillatory spike trains depending on the three parameters, ‘peak rate (*A*)’, ‘baseline (*b*)’ and ‘frequency (*f*)’; see Equation 1. Synchronous outputs of HC and SC neurons are generated with parameters shown below.

### 2.1. Neuron models

All three inhibitory cell types are implemented by neuron models proposed by Hill and Tononi (Hill & Tononi, 2005). The ‘HT’ neuron is a point neuron with simplified Hodgkin–Huxley currents. For reference, we provide a brief review of the neuron model; see (Hill & Tononi, 2005) for details.

The neuronal dynamics obey Equation 2:

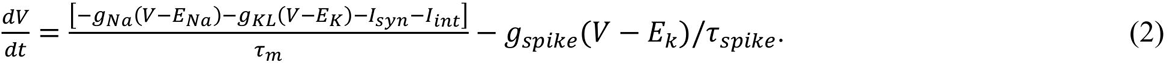

The membrane potentials (V), decayed exponentially with time scale τ_m_, are regulated by sodium (Na) and potassium (K) leak currents with conductance (g_Na_ and g_KL_) and reversal potentials (E_Na_ and E_K_). The fast hyperpolarization current during spikes is simulated with a rectangular spike with a conductance (g_spike_) and decaying time constant (t_spike_). Synaptic events induce dual exponential responses (I_syn_) in the target neurons which are described by rising (τ_1_) and decaying time (τ_2_) constants (Table 2). The reversal potentials for GABA and AMPA are -80 and 0 mV in the model. We did not consider NMDA synapses in the model. The intrinsic ion currents (Iint) are from the original model (Hill & Tononi, 2005).

**Table 2:** Synaptic parameters. All synapses are static and induce double exponential responsesdescribed by τ_1_ and τ_2_. g_peak_ and E_rev_ are the conductance of the synapses and reversal potentials.

| | $g_{\text{peak}}$ | $\tau_1$ | $\tau_2$ | $E_{\text{rev}}$ (mV) |
| --- | --- | --- | --- | --- |
| AMPA | 0.1 | 0.5 | 2.4 | 0 |
| GABA from FS | 0.33 | 1.0 | 7.0 | -80 |
| GABA from non-FS | 0.33 | 1.0 | 20.0 | -80 |

Also, spike threshold (θ) evolves over time with equilibrium (θ_eq_) and time constant (t_θ_), as shown in Equation 2.

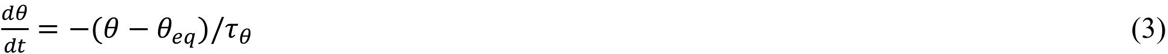

The three cell types have different parameters listed in Table 3. We assume that cholinergic modulation innervates non-FS cells for experimental observations. First, basal forebrain, which provides cholinergic modulation to cortices (Sarter, Parikh, & Howe, 2009), mainly targets somatostatin positive (SST) interneurons (Chen, Sugihara, & Sur, 2015). Second, cholinergic modulation does not modulate the excitability of FS cells (Gulledge, Park, Kawaguchi, & Stuart, 2007). Third, acetylcholine innervates low-threshold spiking interneurons known to express SST via nicotinic receptors but does not modulate the excitability of FS cells (Xiang et al., 1998).

**Table 3:** Neuronal parameters. We list the parameters chosen for the three cell types. g_T_ is the conductance of low-threshold currents (Hill & Tononi, 2005), and the frequency of external background inputs to each cell type is shown in the last column. All cells are implemented using NEST-native neuron models named “ht_neurons” (Gewaltig & Diesmann, 2007), and non-specified parameters are the same as defaults values in NEST.

| | NaP | $t_{\text{spike}}$ | $\theta_{\text{eq}}$ (mV) | $\tau_m$ (msec) | $\tau_\theta$ (msec) | $g_T$ | Ext (Hz) |
| --- | --- | --- | --- | --- | --- | --- | --- |
| Pyr | 1.0 | 2.0 | -51 | 16.0 | 2.0 | 2.0 | 200 |
| FS | 1.0 | 2.0 | -53 | 8.0 | 1.0 | 1.0 | 100 |
| Non-FS | 1.0 | 2.0 | -53 | 8.0 | 1.0 | 2.0 | 100/0 |

### 2.2. Synaptic connections

All synapses in the model have static synaptic weights unlike the depressing synapses in the original model (Hill & Tononi, 2005). FS and non-FS cells provide fast and slowly decaying GABA connections on target neurons, respectively (Traub et al., 2005); 7 msec and 20 msec are chosen for decay time constants for fast and slow kinetics (Table 2). According to the observed pattern (Pfeffer, Xue, He, Huang, & Scanziani, 2013), non-FS cells corresponding to SST cells inhibit FS and Pyr cells, whereas FS cells inhibit FS cells and Pyr cells. These two inhibitory cell types are also consistent with the two functional groups (major regulator and inhibitory selective interneurons) from a recent survey (Jiang et al., 2015). Figure 1A shows the schematic of synaptic connections. When we connect pre-synaptic and post-synaptic populations, we connect cell pairs randomly using connection probabilities (Table 4).

**Table 4:** Connections are randomly generated using the following connectivity. The weights are used to scale synaptic strength (Gewaltig & Diesmann, 2007).

| Connection type | Connection probability | Weights |
| --- | --- | --- |
| External background | N/A | 3.0 |
| Pyr→Pyr | 0.4 | 0.075 |
| Pyr→FS | 0.4 | 0.45 |
| Pyr→non-FS | 0.2 | 0.15 |
| FS→FS | 1.0 | 0.15 |
| FS→Pyr | 0.4 | 0.6 |
| Non-FS→FS | 0.4 | 0.6 |
| Non-FS→Pyr | 0.4 | 0.39 |
| HC to Pyr | 0.2 | 1.2 |
| HC to FS | 0.2 | 0.6 |
| SC to Pyr | 0.2 | 1.2 |
| SC to FS | 0.2 | 1.2 |
| Pyr→downstream | 0.2 | 1.0 |

### 2.3. Simulation of local field potentials

We approximated EEG by calculating local field potentials (LFPs). LFPs were simulated by summing up all the synaptic currents in downstream neurons (Mazzoni, Panzeri, Logothetis, & Brunel, 2008). Then the spectral power density of LFPs was calculated via ‘scipy’ included in python. For each simulation condition, we ran 100 simulations, in which a network is independently instantiated using the same connectivity rule, and reported the average LFP power from them.

### 2.4. Spike-triggered average of LFPs

The coherence between top-down gamma rhythms to A1 and synaptic inputs to downstream neurons was measured with spike-triggered average (STA) of LFPs. In each simulation, we aggregated the 150 msec LFP segments aligned to the spike times of HC cell population which projects top-down gamma rhythms into A1 and averaged them to calculate STA of LFPs. The spectral power of STA of LFPs is calculated in each simulation, and we report the averaged power from 100 independent simulations.

### 2.5. The ratio of firing rate of Pyr cells

In each simulation, we recorded spikes from all Pyr cells and calculated the average firing rate of Pyr cells in the stimulus and baseline (pre-stimulus) periods; the ratio of firing rates between the two periods is also calculated in each simulation. The mean values and standard errors from 100 simulations are reported in Results.

## 3. Results

Figure 1A shows the structure of the model. As seen in the figure, A1 receives afferent inputs from two external populations which model high order cognitive areas (HC) and subcortical areas (SC), respectively. A1 also projects outputs to downstream neurons which model high order auditory areas. In A1, Pyr cells interact with fast-spiking (FS) and non-FS inhibitory interneurons (Fig. 1A); our non-FS cells model low-threshold spiking interneurons which express somatostatin. In the model, non-FS cells inhibit FS and Pyr cells, whereas FS cells inhibit Pyr cells and FS cells. The connectivity pattern is adopted from that which was experimentally reported (Pfeffer et al., 2013). The three cell types, implemented with HT neurons (Hill & Tononi, 2005), exhibit disparate responses to 20 pA tonic currents (Figure 1B); see Methods for details on neuron models. The most active cells are FS cells, and non-FS cells show frequency adaptation, which is consistent with experimental observation (Gibson, Beierlein, & Connors, 1999; Kawaguchi & Kubota, 1997).

In the first subsection (3.1), we discuss the simulation results on the effects of disrupted cholinergic modulation on A1 outputs in the gamma frequency band during baseline (pre-stimulus) and stimulus periods. In the second subsection (3.2), we discuss the potential links between disrupted cholinergic modulation capable of replicating abnormal gamma rhythms and auditory hallucination.

### 3.1. The effects of cholinergic modulation on gamma power generated by A1

We simulate the baseline (0-500 msec) and the stimulus (500-1000 msec) periods. Top-down signals are projected to Pyr and FS cells in A1 (Table 4) during the entire simulation period (1 sec). In contrast, bottom-up signals (i.e., stimulus inputs) are projected to Pyr and FS cells in A1 (Table during the last 500 msec. That is, the two periods are different in terms of the existence of stimulus-inputs.

To simulate 40 Hz auditory-click trains, which were used to estimate stimulus-evoked gamma rhythms in experimental studies (Kwon et al., 1999; Spencer, Salisbury, Shenton, & Mccarley, 2009; Vierling-Claassen et al., 2008), SC neurons fire synchronously at 40 Hz. Similarly, HC neurons, which model high order areas such Par2 (Roopun et al., 2010), fire at 45 Hz. These synchronous inputs are realized by inhomogeneous Poisson events. That is, the firing rate of Poisson spike trains obeys the sinusoidal wave as shown in Equation 1, and the parameters selected for HC and SC neurons are given in Table 1. Importantly, cholinergic modulation depolarizes non-FS cells via cholinergic receptors (Couey et al., 2007; Gulledge et al., 2007; Xiang et al., 1998). Thus, we simulate disrupted cholinergic modulation by reducing the excitability of non-FS cells. Specifically, non-FS cells receive 100 Hz Poisson spike trains in the control condition, whereas those external inputs are removed in the pathological conditions (Table 3).

With synchronous top-down and bottom-up signals (Figure 2A), A1 shows disparate responses between the control and pathological conditions. Non-FS cells fire asynchronously in the control condition (Figure 2B), but they are quiescent in the pathological condition (Figure 2C). FS cell activity appears to be stronger in the pathological condition due to the lack of inhibition from non-FS cells to FS cells. Pyr cell activity also seems stronger in the pathological condition. In both control and pathological conditions, we approximate EEG signals by measuring local field potentials (LFPs) in the downstream neurons receiving afferent inputs from A1. This approximation is based on the observation that synaptic currents onto Pyr cells are the dominant factor for both LFPs and EEG (Destexhe & Bedard, 2013). Specifically, synaptic currents into the downstream neurons are accumulated over time (Mazzoni et al., 2008); see Methods.

**Figure 2:**
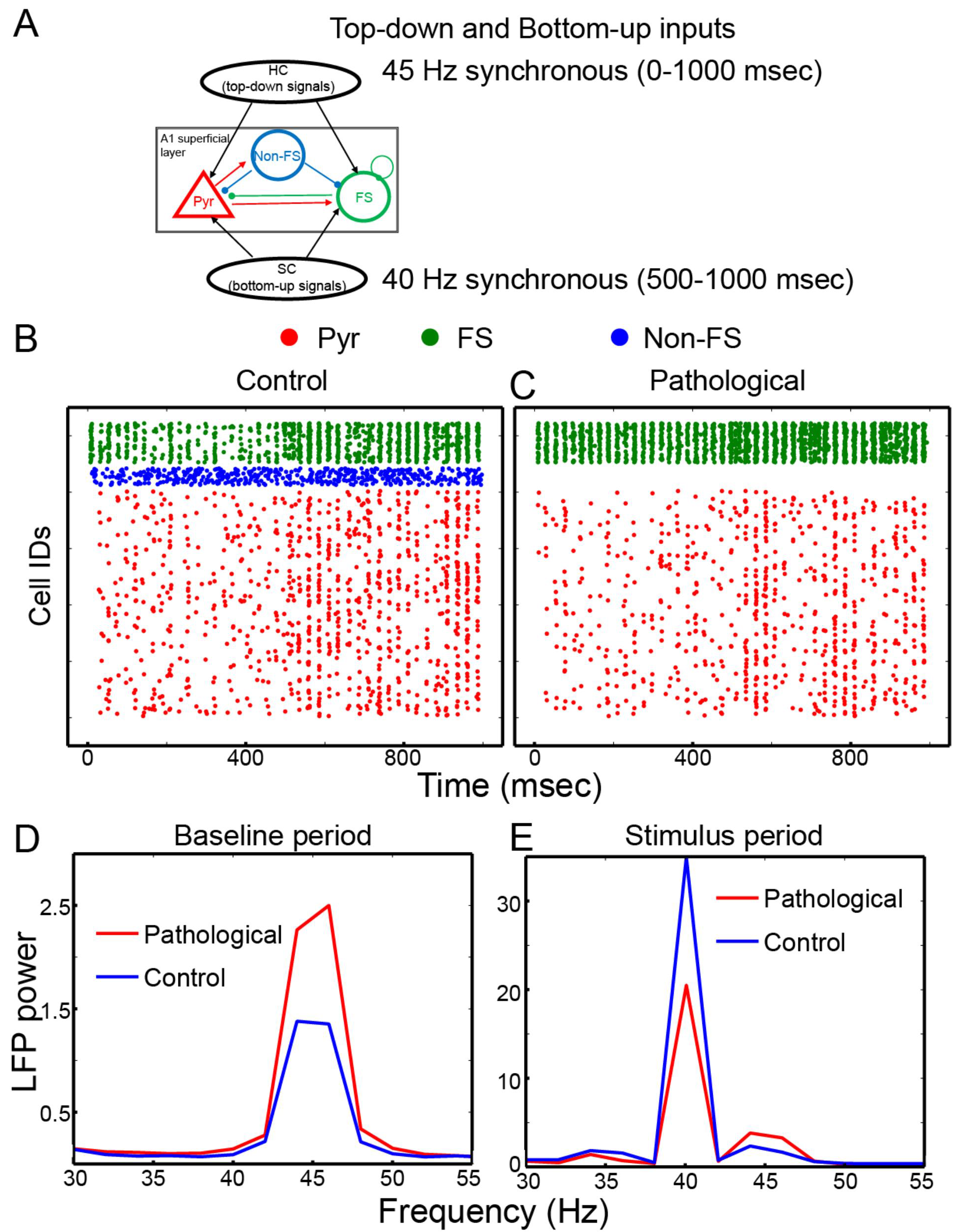
The network responses depending on the excitability of non-FS cells. **(A)**, Theschematics of external inputs to A1. **(B)**, The red, blue and green dots represent action potentials of Pyr, non-FS and FS cells in the control condition. Each row in the y-axis is the ids of cells. **(C)**, Action potentials in the pathological condition. **(D)**, The comparison of the spectral power of LFPs in the baseline period between the control and pathological conditions; the scale of LFP power is arbitrary. **(E)**, The comparison of LFP power between the two conditions.

In the baseline period (prior to bottom-up inputs from SC neurons), 45 Hz rhythms are induced by top-down gamma rhythms. The induced peak power is higher in the pathological condition than in the control condition (Figure 2D), which is consistent with the enhanced baseline gamma rhythms (Hirano et al., 2015; Spencer, 2011). We note that Pyr cells fire more strongly but more asynchronously in the baseline period when non-FS cells are active. This accounts for the weaker baseline gamma-band power in the control condition (Figure 2D). In the stimulus period, 45 Hz rhythms are reduced, and 40 Hz rhythms induced by bottom-up signals are generated (Figure 2E). That is, A1 responds to bottom-up gamma rhythms rather than to top-down gamma rhythms in the stimulus period. More importantly, as seen in Figure 2E, the peak power at 40 Hz is higher in the control condition, which is consistent with reduced stimulus-evoked gamma rhythms (Kwon et al., 1999; Lee, Whittington, & Kopell, 2015; Vierling-Claassen et al., 2008).

#### 3.1.1. Non-FS cells indirectly regulate Pyr cell activity

Simulation results above support that disrupted cholinergic modulation can underlie abnormal gamma rhythms. In the pathological condition, non-FS cells became quiescent, indicating that the inhibition of non-FS cells underlies abnormal gamma rhythms observed in people with schizophrenia. To further study gamma rhythms’ modulation by the inhibition of non-FS cells, we repeat the same simulations but varied the strengths of non-FS cells impinging onto FS or Pyr cells. Figure 3 shows the spectral power of LFPs depending on the strengths of the inhibition of non-FS cells. We note two germane observations. First, the baseline LFP power in the baseline period decreases, as non-FS-Pyr cell connections are strengthened (Figure 3A). Second, the strength of inhibition onto FS cells is positively correlated with LFP power in the stimulus period (Figure 3B). That is, non-FS cells play distinctive roles in the baseline and stimulus periods. In the baseline period, FS cells reduce A1’s responses to top-down gamma rhythms by projecting inhibition onto Pyr cells. In contrast, during the stimulus period, they increase Pyr cell responses to stimulus inputs.

**Figure 3:**
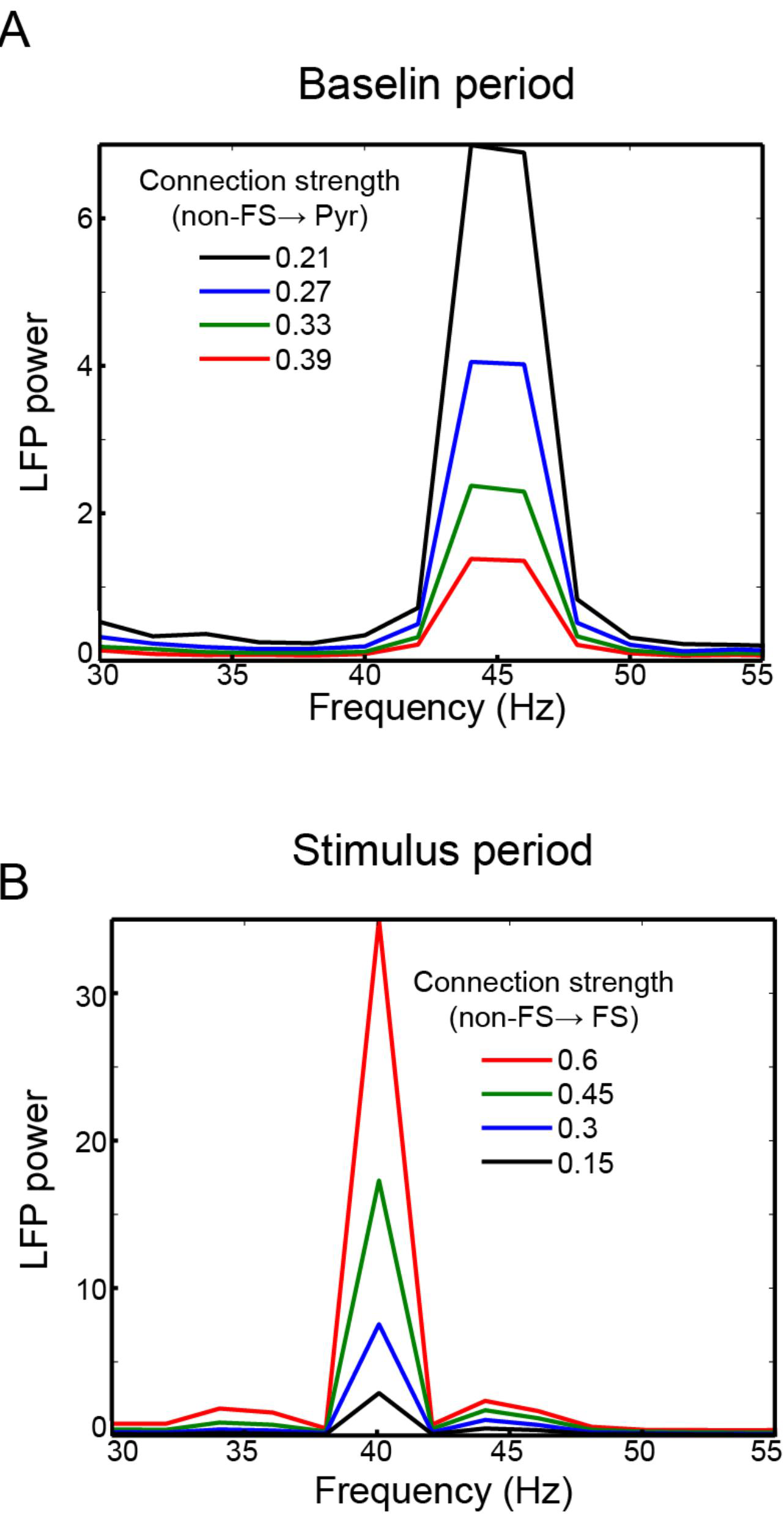
The effects of inhibition of non-FS cells. **(A)**, The modulation LFP power via inhibition from non-FS to FS cells in the baseline period. **(B)**, The modulation of LFP power via inhibition from non-FS to Pyr cells in the stimulus period.

#### 3.1.2. Top-down signals can enhance A1 responses to auditory stimuli

In the model, baseline gamma rhythms are induced by top-down gamma rhythms innervating both Pyr and FS cells (Figure 1A and Table 4). As top-down gamma rhythms target FS cells preferentially, they may suppress A1 responses. If noise-driven responses are suppressed, stimulus-evoked responses may be easily recognized; that is, the signal-to-noise ratio of A1 responses may become higher. We further study this possibility by calculating the ratio of stimulus-evoked responses to the baseline responses, which is referred to as SNR hereafter. In this simulation condition, we assume that SC neurons fire asynchronously at 60 Hz to consider more generic stimulation condition (Figure 4A); A1 cells fire asynchronously in response to synchronous inputs at higher frequency than 40 Hz (Wang, 2003). Examples of responses of the model A1 in the control and pathological conditions are displayed in Figure 4B and C, respectively. As expected, stimulus-evoked Pyr cell responses are stronger in the control period. To quantify how effectively top-down gamma rhythms enhance SNR, we estimate SNR depending on the peak firing rate of HC cells (Method). We run 100 independent simulations, in each of which SNR is calculated (Method). Figure 4D shows that the mean value and standard errors of SNR values from 100 simulations. As seen in the figure, SNR increases, as the peak firing rate of top-down signals increases, confirming that top-down gamma rhythms make A1 generate more distinct responses, when compared to baseline activity.

**Figure 4:**
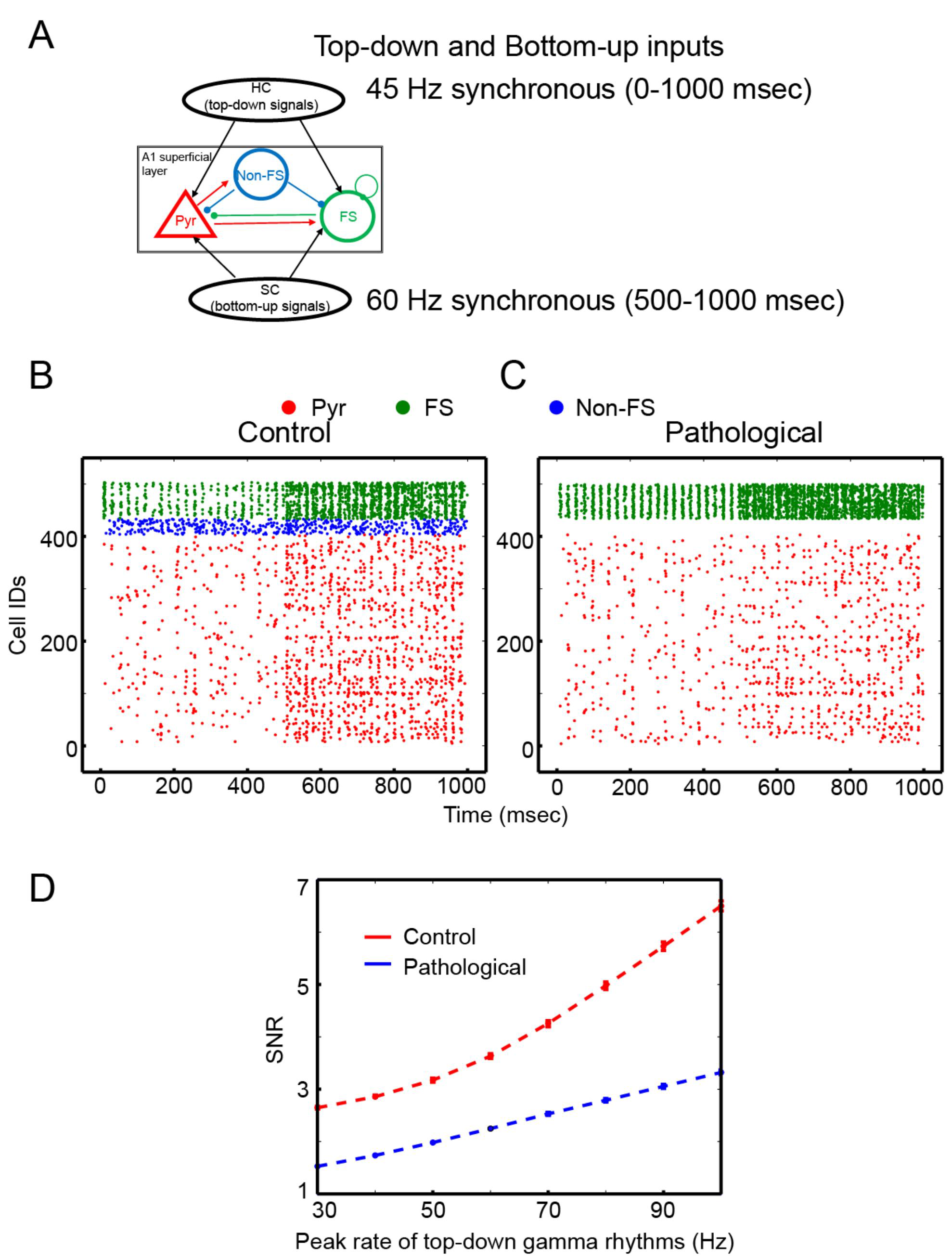
The functional roles of top-down gamma rhythms. **(A)**, The schematics of external inputs to A1. **(B)** and **(C)**, Spikes generated by three cell types in the control and pathological conditions, respectively. **(D)**, SNR depending on the peak rate of top-down gamma rhythms (i.e., *A* in Equation 1).

### 3.2. The potential pathology of auditory hallucination

Our simulation results suggest that two inhibitory cell types (FS and non-FS cells) play distinct roles in auditory signal processing. FS cells make A1 responses sharper, whereas non-FS cells make A1 responses more reliable. Then, do these cell types contribute to auditory hallucination? If so, what are their functional roles? Below, we describe simulations regarding potential pathophysiology of auditory hallucination.

#### 3.2.1. Top-down gamma rhythms can entrain A1 Pyr cells in the pathological condition

In the model, top-down gamma rhythms enhance signal-to-noise ratio of A1 responses by reducing noise-driven responses. However, the enhanced baseline gamma rhythms suggest that A1 Pyr cells can fire in response to top-down signals in the pathological condition. We further examine how effectively individual spikes of HC cells can induce synaptic inputs from A1 to downstream neurons. In doing so, we remove the stimulus (Figure 5A) and calculate the spike-triggered average (STA) of LFPs; see Methods. When HC neuron spikes reliably induce A1 to project inputs to downstream neurons, STA of LFPs will become oscillatory. Otherwise, the oscillatory responses will be feeble. The spectral power of STA of LFPs is stronger (Figure 5A) in the pathological condition than in the control condition, confirming that top-down gamma rhythms induce A1 to generate outputs despite the absence of auditory stimulus. Such synchronous outputs may be critical in mediating interareal communications (Fries, 2005).

**Figure 5:**
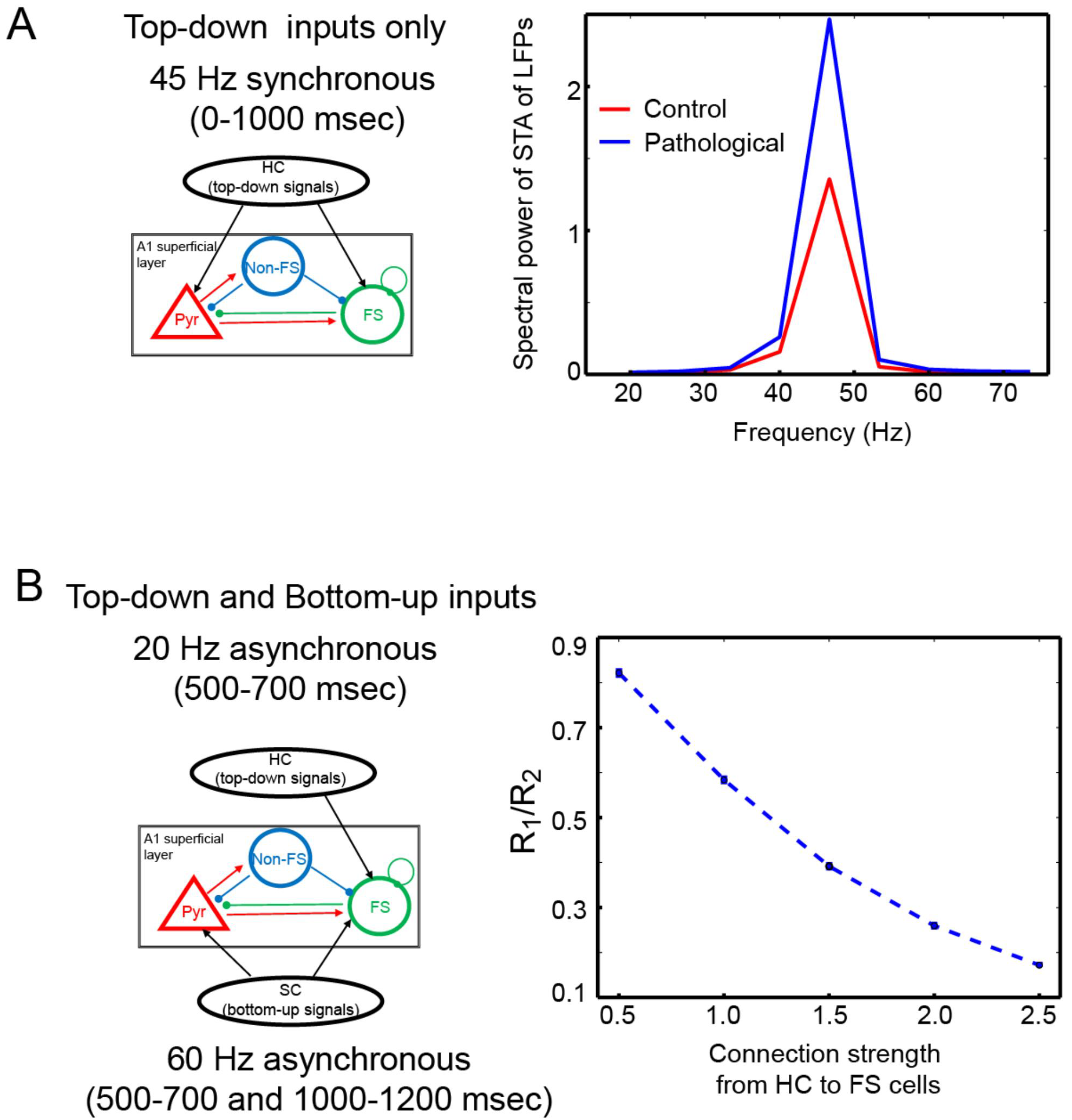
Incorrect A1 responses. **(A)**, The spectral power of STA of LFP using spikes of HC cells (i.e., top-down gamma rhythms). **(B)**, The comparison of Pyr cell responses between the two stimulus periods (500-700 and 1000-1200 msec). The firing rates (R_1_ and R_2_) of Pyr cells are calculated during the former and latter periods, respectively, and the ratio R_1_/R_2_ is estimated in each simulation. The mean values and standard errors from 100 simulations are shown in the right panel.

#### 3.2.2. Asynchronous top-down signals can subserve corollary discharge

A1 responses to self-generated sound are reduced (Ford & Mathalon, 2005; Schneider, Nelson, & Moony, 2014), and the corollary discharge, a copy of motor signals projected to A1, has been proposed as an underlying mechanism. The simulation results above suggest that FS cells may mediate such corollary discharge. To address this assertion, we estimate A1 responses with and without asynchronous top-down signals to FS cells; as shown in Figure 5B, these asynchronous inputs are introduced to FS cells only. Specifically, two stimulus inputs are introduced to A1 during 500-700 msec and 1000-1200 msec. In this simulation condition, for the sake of generality, both top-down and bottom-up signals are assumed to be asynchronous (Figure 5B); 20 Hz and 60 Hz Poisson spike trains are generated by HC and SC cells, respectively. We assume that the former inputs are induced by self-generated sound and provide top-down signals to FS cells only during 500-700 msec (Figure 5B). If FS cells could subserve corollary discharge, Pyr cell responses to the former inputs would be lower than those to the latter inputs. With this assumption in mind, we compare Pyr cell response to the former and the latter bottom-up inputs. In each simulation, we calculate the average Pyr activity (R1, and R2) in two stimulus periods (500-700 and 1000-1200 msec), respectively. The ratio of R1 to R2 is calculated, which is dependent on the strength of connections from HC to FS cells (Figure 5B). We repeat 100 simulations and calculate the mean and standard errors from them. As expected, Pyr cell responses are reduced when top-down signals innervated FS cells. In addition, the reduction of Pyr responses to the (simulated) self-generated sound is more prominent, as the strength of connections from HC neurons to FS cells increases (Figure 5B).

## 4. Discussion

We utilized a computational model of A1 to elucidate the potential pathophysiology by which abnormal gamma rhythms are induced in schizophrenia. Our simulation results support our hypothesis that disrupted cholinergic modulation underlies the pathophysiology of schizophrenia. Below we discuss the implications of our simulation results.

### 4.1. FS cell-mediated corollary discharge

Corollary discharge failure is one of the leading theories of auditory hallucination. It has been reported that the functional connectivity between frontal lobe that generate corollary discharge and temporal lobe decreased in people who experience auditory hallucination (Ford & Mathalon, 2005), raising the possibility that corollary discharge from language areas can be weakened in people with schizophrenia. Our simulation results show that asynchronous top-down signals to FS cells suppress A1 responses and that the reduction of A1 responses to a (simulated) self-generated sound is weaker when the connection strength from HC to FS cells is weaker. Therefore, we propose that FS cells mediate corollary discharge.

Could non-FS cells also mediate corollary discharge? We argue against the possibility that non-FS cells can be mediators for corollary discharge for two reasons. First, non-FS cells show frequency adaptation, and thus the inhibition of non-FS cells onto Pyr cells may be weakened over time, and Pyr cells may respond to a long-lasting self-generated sound. Second, the third class of inhibitory cells (Rudy et al., 2011), vasoactive intestinal peptide positive (VIP) cells, inhibits SST cells (Pfeffer et al., 2013) which correspond to non-FS cells in the model. Since cingulate cortex innervates VIP cells most strongly (Zhang et al., 2014), non-FS cell activity may be suppressed during top-down process involving the cingulate area. That is, when cingulate cortex activates VIP cells, corollary discharge may be disabled if non-FS cells mediate corollary discharge. These properties of non-FS cells suggest that FS cells are more appropriate for mediating corollary discharge.

### 4.2. False perception induced by top-down gamma rhythms

Top-down gamma rhythms in our model make A1 responses stronger, relative to noise-driven responses. This is consistent with the notion that attention enhances auditory responses selectively as search light does (Fritz et al., 2007). Together with the observation that FS cell activity is enhanced during attention (Mitchell, Sundberg, & Reynolds, 2007), we propose that top-down gamma rhythms can contribute to attentional gain control. More importantly, in the model, top-down gamma rhythms can induce A1 to send stronger synchronous synaptic inputs to its target areas in the pathological condition than in the control condition. According to the ‘communication through coherence hypothesis’ that synchronous activity subserve interareal communication (Fries, 2005), such abnormally strong synchronous synaptic inputs to downstream neurons can be perceived as actual auditory signals, which induces auditory hallucination (Kristiina Kompus et al., 2013; Powers III et al., 2016; F. A. Waters et al., 2006). Moreover, cortical rhythms are generated during speech processing and sensory prediction (Arnal, Wyart, & Giraud, 2011; Giraud & Poeppel, 2012), and thus our model proposes potential mechanisms by which top-down signals mediating expectation or speech information penetrate perceptual system and induce hallucination. This could explain why auditory hallucination often includes verbal hallucination. In addition, our simulation results suggest that A1 is activated during auditory hallucination and reveals the difficulty of distinguishing auditory hallucination from real stimuli in some cases; A1 activation will certainly induce vivid hallucination.

### 4.3. Broadband peak power increase induced by narrowband top-down gamma rhythms

In the model, baseline gamma rhythms show narrowband peak power around 45 Hz. In the experiments, however, the enhanced broadband power was observed (Hirano et al., 2015). The origins of broadband gamma rhythms remain unclear, while narrowband gamma rhythms can be explained by the interplay between pyramidal cells and interneurons or recurrent inhibition among interneurons (Whittington, Traub, Kopell, Ermentrout, & Buhl, 2000). A theoretical study proposed that broadband gamma rhythms can be generated by interactions among multiple oscillators, which is called ‘synchronous chaos’ (Battaglia, Brunel, & Hansel, 2007), and interactions between laminar layers, each of which generates narrowband oscillatory activity, were shown to induce broadband gamma rhythms elicited by visual stimuli (Battaglia & Hansel, 2011). That is, narrowband top-down gamma rhythms may be capable of triggering broadband rhythms via interlaminar interactions. Similarly, when top-down gamma rhythms induce narrowband oscillatory activity in a single cell assembly, the neighboring assemblies may also generate rhythmic activity via lateral excitation. As multiple assemblies are stimulated, the interactions among them will generate broadband rhythms via synchronous chaos.

### 4.4. Correlation between abnormal gamma rhythms and auditory hallucination

Our simulation results suggest that top-down gamma rhythms induce A1 to bring about false perception, predicting that baseline gamma rhythm enhancement may be positively correlated to auditory hallucination. However, an earlier experimental study (Spencer, 2011) did not find any correlation. This may be attributed to the fact that subjects in the study participated in passive listening tasks, in which top-down influence may not be strong. That is, top-down gamma rhythms may be strong enough to enhance baseline power but too weak to induce sufficient false outputs to generate auditory hallucination.

Interestingly, the gamma band power in the stimulus period was reported to be positively correlated with the extent of hallucination (C Mulert, Kirsch, Pascual-marqui, Mccarley, & Spencer, 2011; Spencer, Niznikiewicz, Nestor, Shenton, & McCarley, 2009). The gamma band power in the stimulus period, however, can also depend on the effective baseline activity (Flynn et al., 2008). As auditory stimuli innervate a subset of auditory neurons, most auditory neurons remain in the effective baseline period. As long as top-down gamma rhythms are projected to the auditory neurons, which are not directly stimulated by sound, they produce gamma rhythms and contribute to the total gamma band power during stimulus presentation. Our simulation results predict that the baseline power induced by the auditory neurons receiving only top-down gamma rhythms will be stronger in the pathological condition, suggesting that the total power in the stimulus period is also enhanced in the pathological condition. This accounts for the positive correlation between gamma band power and auditory hallucination score (C Mulert et al., 2011; Spencer, Niznikiewicz, et al., 2009).

### 4.5. Limits of our model and future direction

We study cholinergic modulation of A1 responses to bottom-up and top-down signals and its potential contribution to pathophysiology of auditory hallucination, and for this study, other phenomena related to schizophrenia and auditory hallucination are ignored.

First, we focused on the effects of reduced non-FS cell activity, as non-FS cells’ excitability is regulated directly by cholinergic modulation (Couey et al., 2007; Xiang et al., 1998). Our simulation results support the importance of cholinergic modulation in regulating gamma rhythms in the auditory system. However, this does not exclude the possibility that FS cells can be directly involved with aberrant gamma rhythms or auditory hallucination. In fact, the hypofunction of N-methyl-D-aspartate (NMDA) receptors and the reduction of excitatory inputs in FS cells and FS cell outputs have been thought to underlie the pathophysiology of schizophrenia (Jadi, Margarita Behrens, & Sejnowski, 2015; Pittman-Polletta, Kocsis, Vijayan, Whittington, & Kopell, 2015; Spencer, 2009).

Second, in the auditory system, gamma rhythms are modulated by low-frequency rhythms such as delta-and theta rhythms (Lakatos et al., 2010; Lakatos, Chen, O’Connell, Mills, & Schroeder, 2007; Tsunada, Baker, Christison-Lagay, Davis, & Cohen, 2011), which was referred to as ‘oscillatory hierarchy’ (Lakatos et al., 2005). Interestingly, these low-frequency rhythms were also abnormal in schizophrenia. In this study, we focused on investigating the effects of cholinergic modulation on gamma rhythms and ignored the interactions between gamma rhythms and low frequency rhythms. For our next study, we will study the underlying mechanisms of abnormalities in low-frequency rhythms.

In the future, we plan to extend this research to incorporate those additional complexities in two ways. First, we should probe the abnormalities of FS cells in schizophrenia and their interactions with disrupted cholinergic modulation. Second, considering the ‘dysconnectivity’ hypothesis that their pathophysiology involves aberrant interactions among multiple brain regions, we will build an extended multiple-area model to study the aberrant interactions. Such a multi-brain region model would help us address abnormal low-frequency rhythms, as theta rhythms are prominent in the hippocampus.

